# The effects of seasonal climate variability on dengue annual incidence in Hong Kong: A modelling study

**DOI:** 10.1101/2020.01.20.912097

**Authors:** Hsiang-Yu Yuan, Jingbo Liang, Pei-Sheng Lin, Kathleen Sucipto, Mesfin Mengesha Tsegaye, Tzai-Hung Wen, Susanne Pfeiffer, Dirk Pfeiffer

## Abstract

In recent years, dengue has been rapidly spreading and growing in the tropics and subtropics. Located in southern China, Hong Kong’s subtropical monsoon climate may favour dengue vector populations and increase the chance of disease transmissions during the rainy summer season. An increase in local dengue incidence has been observed in Hong Kong ever since the first case in 2002, with an outbreak reaching historically high case numbers in 2018. However, the effects of seasonal climate variability on recent outbreaks are unknown. As the local cases were found to be spatially clustered, we developed a Poisson generalized linear mixed model using pre-summer monthly total rainfall and mean temperature to predict annual dengue incidence (the majority of local cases occur during or after the summer months), over the period 2002-2018 in three pre-defined areas of Hong Kong. Using leave-one-out cross-validation, 5 out of 6 observations of area-specific outbreaks during the major outbreak years 2002 and 2018 were able to be predicted. 42 out of a total of 51 observations (82.4%) were within the 95% confidence interval of the annual incidence predicted by our model. Our study found that the rainfall before and during the East Asian monsoon (pre-summer) rainy season is negatively correlated with the annual incidence in Hong Kong while the temperature is positively correlated. Hence, as mosquito control measures in Hong Kong are intensified mainly when heavy rainfalls occur during or close to summer, our study suggests that a lower-than-average intensity of pre-summer rainfall should also be taken into account as an indicator of increased dengue risk.

## Introduction

Dengue is one of the most prevalent vector-borne arboviruses, transmitted between humans by *Aedes* mosquitoes. The number of dengue cases reported to the World Health Organization (WHO) has increased steadily from an average of less than a thousand cases per year globally in the 1950s to over 3.34 million in 2016^1, 2^. Moreover, a recent study estimated that 390 million dengue infections occur each year^3^ while another indicated that 3.9 billion people in 128 countries are at risk of infection^4^. The number of cases is still increasing as the disease spreads to new areas and explosive outbreaks occur. This global expansion of dengue virus (DENV) can be due to several causes, including environmental and climate factors. Climate is an important determinant of vector-borne disease epidemics as it directly influences the abundance and distribution of the vector. Together with environmental factors such as the degree of urbanisation, climate variations may result in the geographic expansion of mosquitoes and dengue into new areas^5^.

Studies have shown that a positive association exists between temperature and dengue transmission^6–9^. One possible reason is that warmer temperatures lead to a higher abundance of *Aedes* mosquitoes by increasing their survival and development rates^10, 11^. Furthermore, the extrinsic incubation period (EIP; the time required for viruses to become transmissible after the initial infection of a mosquito) of DENV may be shortened under warmer conditions^12, 13^, resulting in increased efficiency of viral transmission by allowing more mosquitoes to transmit the virus after exposure. The effect of rainfall amount on dengue incidence, on the other hand, is more controversial. Rainfall has been generally thought to be a positive predictor of dengue vector abundance as it provides essential mosquito habitats^11, 14^. Recent studies, however, have demonstrated that excess rain may also flush away eggs, larvae or pupae of mosquitoes and remove mosquito breeding sites^15, 16^, hence negatively effecting dengue incidence. Another study has found that dry seasons followed by a period of excess rainfall can lead to a high risk of dengue outbreaks, indicating that either positive or negative effects can occur, however, with different time lags^17^. Therefore, a better understanding of how rainfall affects dengue infection rates in areas with large outbreaks or increasing incidence could provide better projections of climate effects on dengue expansion.

The South-East Asia and Western Pacific regions, two of the six WHO regions, together have accounted for nearly 75% of the current global burden of dengue infections^1^. A rapid increase of dengue incidence has been recently observed near the centre part of the Western Pacific region, such as in Guangzhou (the capital city of Guangdong province in southern China) and southern Taiwan^18, 19^. Hong Kong, located geographically close to both Taiwan and Guangzhou, has a typical East Asian subtropical monsoon climate, characterised by higher temperatures and copious amounts of rain in the spring and summer months^20^, which may favour dengue vector abundance and cause outbreaks later during or after summer. Despite the successful dengue control measures introduced after the first local case was identified in Hong Kong in 2002^21^, there has been a gradual increase in dengue incidence after 2014 with the highest number of dengue so far recorded in 2018. Identifying the role of climate variables, particularly the effects of rainfall, on recent distribution trends in Hong Kong is urgently needed in order to understand the recent large outbreak and geographical expansion of dengue in the Western Pacific region or globally. However, the effects of pre-summer rainfall on dengue incidence around southern China are often ignored since the rainfall season normally has a lead time up to three to five months before the outbreaks. These lag effects were often not considered in many of the previous studies in this region^22–25^.

This study aims to enable prediction the annual number of dengue cases using seasonal monthly temperature and rainfall data before summer and assessing the effects of climate variability on the recent increase of dengue incidence in Hong Kong. Additionally, given that all dengue outbreaks/infections in Hong Kong occurred as either local clusters or sporadic cases in certain areas instead of large regional scale epidemics, the study utilised a generalized linear mixed model that contains area-specific random effects to account for environmental factors which are difficult to measure and fixed effects to account for climate effects on dengue transmission. Understanding how the climate may affect annual dengue incidence is important in order to forecast future possible dengue outbreaks, thereby allowing for more effective mosquito control measures in each region, leading to early preparedness and prevention of dengue.

## Results

### Area-Specific Dengue Cases in Hong Kong

After the first local case of dengue fever was discovered in 2002, a large dengue outbreak occurred in the same year and another in 2018 while sporadic cases mainly occurred between 2014 and 2017. We divided Hong Kong into 3 areas: i) New Territories-South (NTS), ii) New Territories-North (NTN), and iii) Hong Kong Island & Kowloon (HKL), based on the administrative districts, the population density, and the usage of the Mass Transit Railways (see Methods). The existence of spatial heterogeneity of dengue infections was observed between the defined areas (Fig. 1 and its inset). For example, HKL and NTS together consisted of all the infections during 2018 outbreak, while HKL and NTN contained all the sporadic cases from 2014 to 2017. Note the special case of the year 2002 when 17 out of 20 locally acquired dengue cases, which included one case transmitted via blood transfusion, were all located around a construction site on an island geographically at the centre of the three areas (shown in red in Fig. 1). After the blood transfusion case was removed, the remaining cases were equally assigned to the three areas with one extra case assigned to NTN.

**Figure 1.**
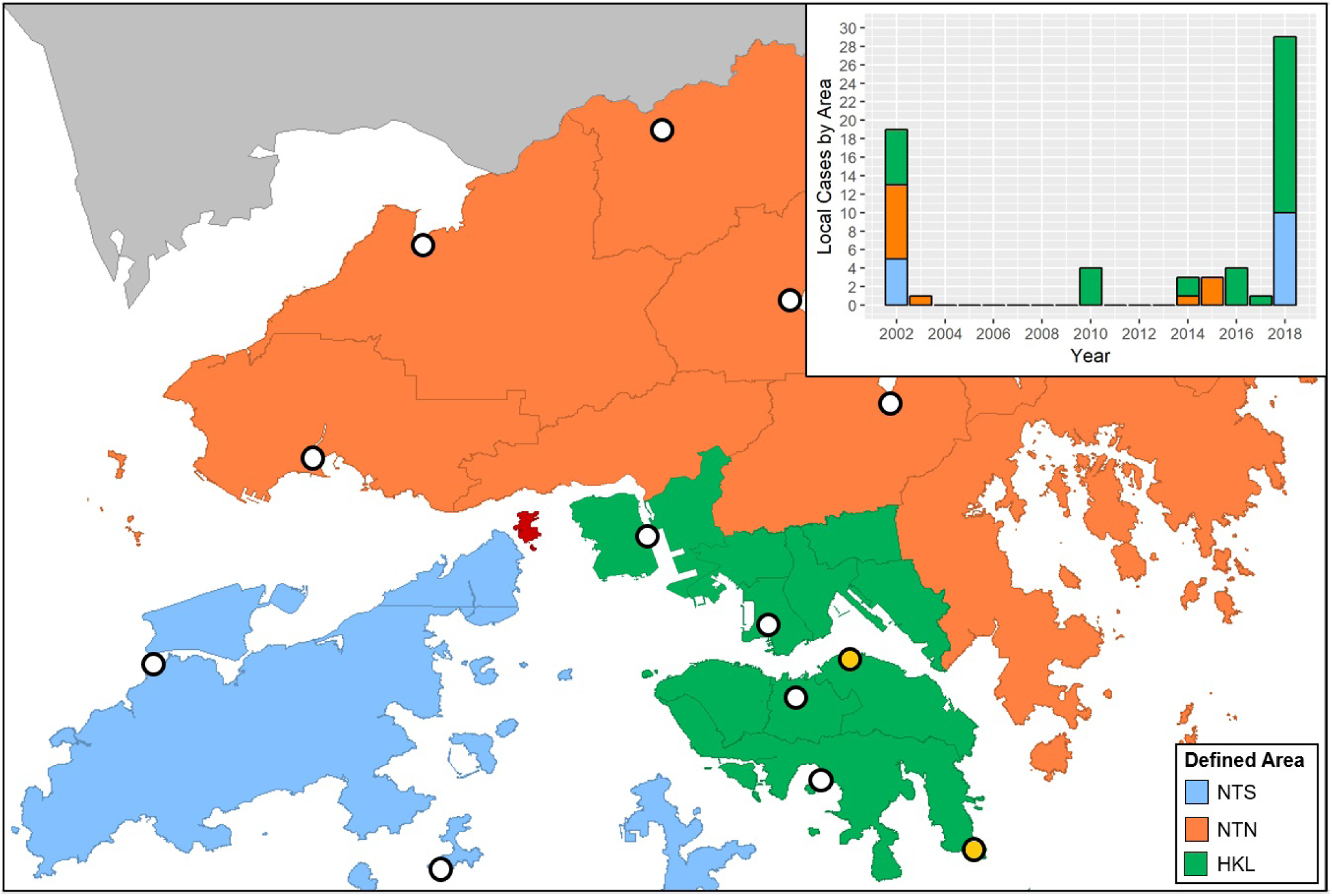
The area division: New Territories - South (NTS; light blue), New Territories - North (NTN; orange), and Hong Kong & Kowloon (HKL; green) along with the selected automatic weather stations (white circles) and rainfall stations (light yellow circles). The grey lines indicate the official district borders. As Ma Wan Island (red) is located at the center, all the 17 non-blood transfusion cases on that island are assigned into the 3 areas in our study: 5 cases in NTS, 6 cases in NTN, and 5 cases in HKL. The inset provides annual local case numbers from 2002 to 2018

### Climate Predictors Selection

In order to select suitable climate parameters for area-specific annual dengue forecasting, we compared 4 models with different combinations of temperature (either monthly mean or minimum; indicated by *T*_*mean*_ or *T*_*min*_) and rainfall (either monthly total or maximum; indicated by *R*_*tot*_ or *R*_*max*_). These climate parameters were suggested by the prior studies conducted in nearby countries^8, 26, 27^, but their association with recent dengue outbreaks in Hong Kong is still unknown and the effects of rainfall on dengue spreading are mixed. Furthermore, spatial differences in the climate data were not considered when the effects of monthly climate predictors were analysed^8^. Thus, all the climate data within the studied years from 2002 to 2018 were retrieved from 11 weather stations and 2 automatic rainfall stations across the defined areas (white and yellow circles Fig. 1), with the averages from January to August shown in Fig. 2. In general, the highest temperatures and the highest rainfalls were recorded between June and August and after April, respectively.

**Figure 2.**
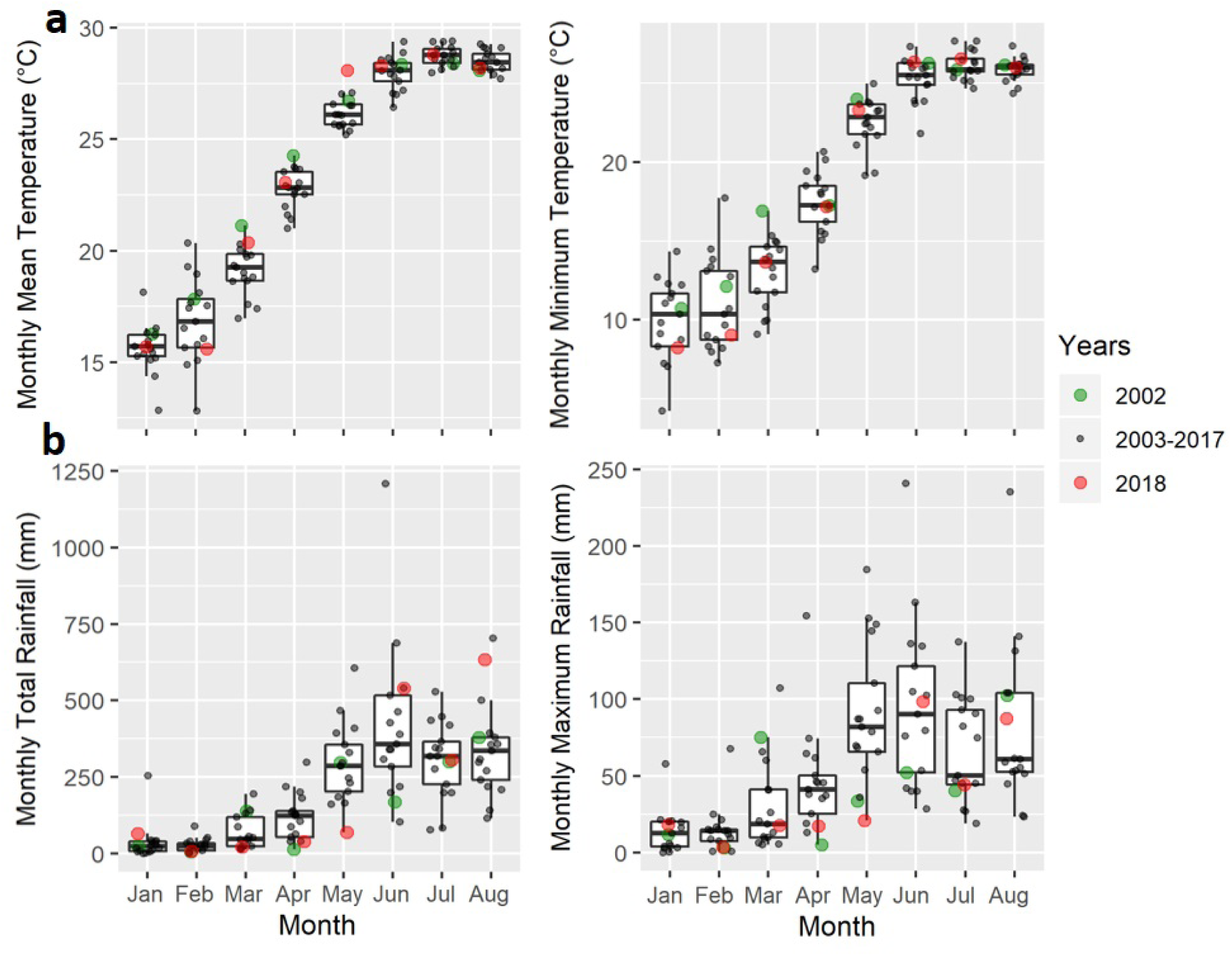
The monthly climate data for the whole of Hong Kong from 2002 to 2018. (a) Monthly mean and monthly minimum temperature, and (b) monthly total and monthly maximum rainfall from January to August for the years 2002, 2003-2017 combined, and 2018 are shown. Climate data were retrieved from 11 weather stations and 2 automatic rainfall stations across the three pre-defined areas

Since most of the local (indigenous) dengue cases occurred during the summer seasons starting in or after August (Fig. 3) and in Hong Kong mosquitoes are not commonly observed during the winter until March^28, 29^, for each combination of climate parameters we used monthly climate predictors (Fig. 4) from March to August in the 3 defined areas to forecast area-specific annual dengue cases using a Poisson mixed effects model. The predictors were first normalised in order to compare the effects between each predictor. The stepwise algorithm based on the Akaike Information Criterion with a correction (AICc) was used to select the monthly predictors for each model (Supplementary Table S1). The best-fitting model was determined according to both the AICc and Bayesian Information Criterion (BIC).

**Figure 3.**
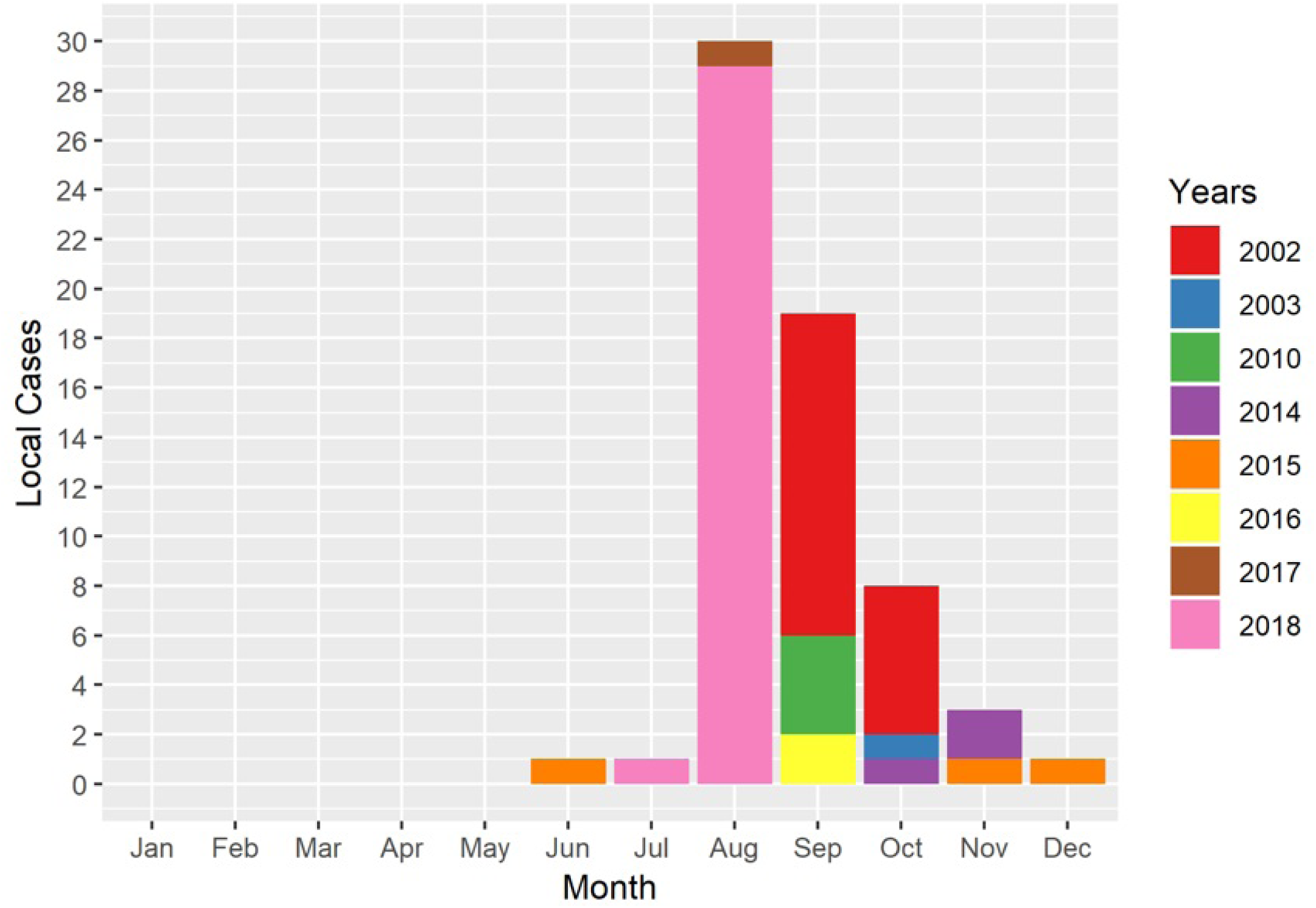
The number of monthly reported local dengue cases from 2002 to 2018. Different colours represent the number of dengue cases in different years.

**Figure 4.**
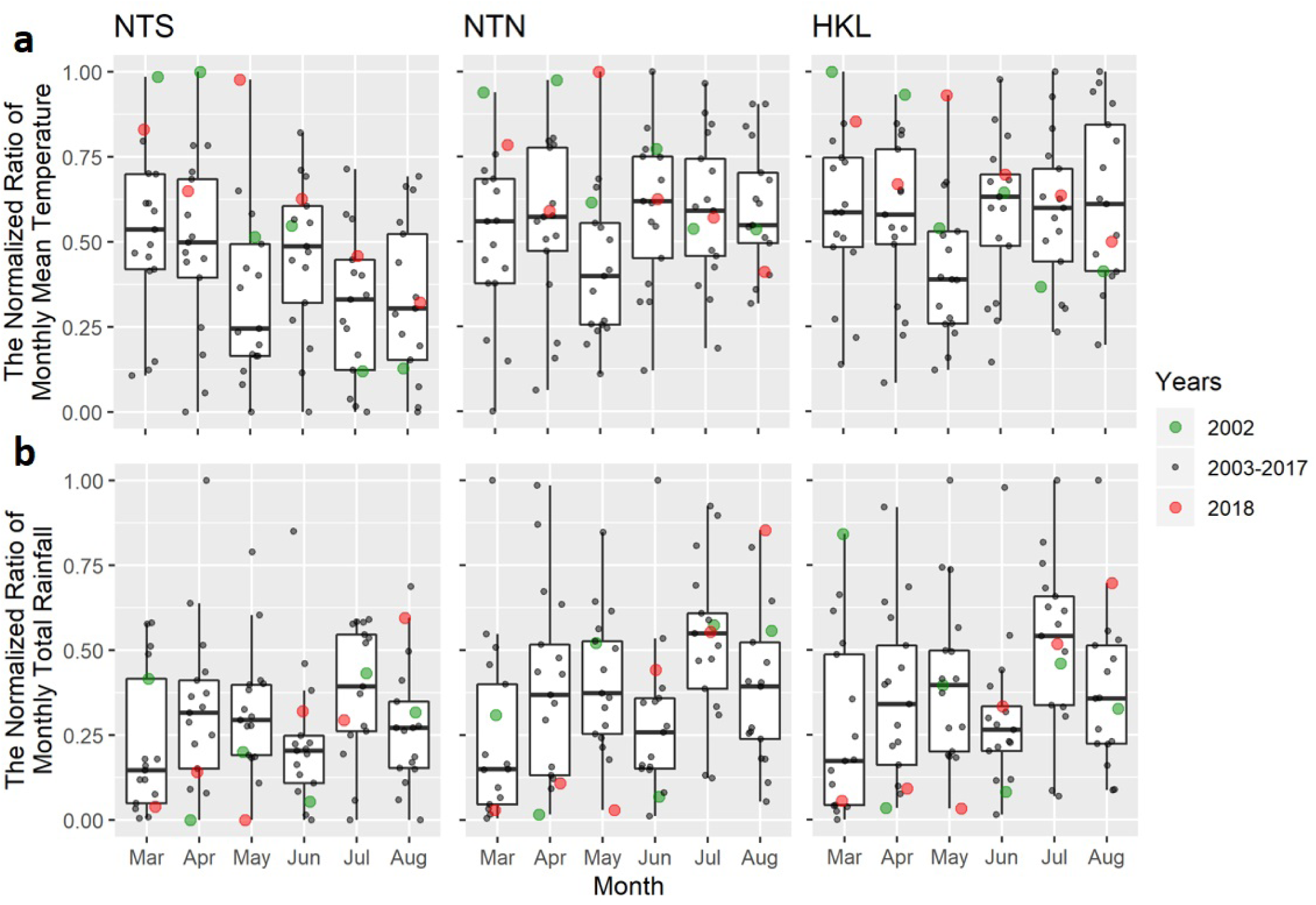
The normalised climate predictors for each area using (a) the monthly mean temperature and (b) the monthly total rainfall. The areas NTS, NTN, and HKL are defined as in Figure 1.

### Area-specific Annual Dengue Forecasting

The results showed that the models with the *T*_*mean*_ climate parameters set performed better than the models with the *T*_*min*_ parameters (Table 1). The best-fitting model, Model: *T*_*mean*_ + *R*_*tot*_ was used as our predictive model for dengue cases while Model: *T*_*mean*_ + *R*_*max*_ was used as an alternative model as both the Δ*AICc* and Δ*BIC* results were equal to only 1 for Model: *T*_*mean*_ + *R*_*max*_. The best-fitting model estimated using the Laplace approximation is given by:

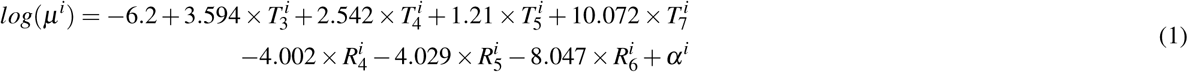

where the standard deviation of the random intercepts *σ* = 0.789. A significance test was performed for each fixed effect predictor using the likelihood ratio test. P-values were less than 0.05 for 8 out of 9 regression coefficients (Table 2). The random intercepts represent different levels of effects resulting from unobserved area-specific factors, which may include varying land usages or population densities between different areas.

**Table 1.** The AICc and BIC of each model with the selected predictors.The most optimal set of predictors was determined using the StepAICc for each climate parameters set. Model: *T*_*mean*_ + *R*_*tot*_ was selected as the best-fitting model and Model: *T*_*mean*_ + *R*_*max*_ was used as the alternative model.

| Set | $T_{mean}$ | | $T_{min}$ | |
| --- | --- | --- | --- | --- |
| | $R_{tot}$ | $R_{max}$ | $R_{tot}$ | $R_{max}$ |
| $AICc$ | 107.84 | 108.84 | 134.34 | 117.91 |
| $\Delta AICc$ | 0.00 | 1.00 | 26.50 | 10.07 |
| $BIC$ | 120.83 | 121.83 | 147.33 | 132.39 |
| $\Delta BIC$ | 0.00 | 1.00 | 26.50 | 11.56 |

**Table 2.**
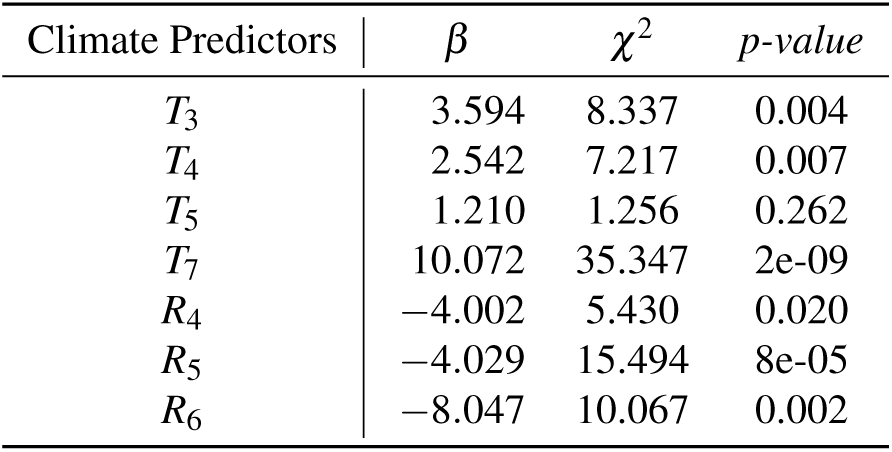
The Likelihood Ratio Test (LRT) results for each selected climate predictor in the best-fitting model. Each predictor was removed and compared to the original model using ANOVA. The p-value was then calculated given that the test statistic of the LRT is asymptotically approximating a *χ*^2^ distribution. *T*_*m*_ and *R*_*m*_ denote the temperature and rainfall predictors in the *m*^*th*^ month of the year.

| Climate Predictors | $\beta$ | $\chi^2$ | $p$-value |
| --- | --- | --- | --- |
| $T_3$ | 3.594 | 8.337 | 0.004 |
| $T_4$ | 2.542 | 7.217 | 0.007 |
| $T_5$ | 1.210 | 1.256 | 0.262 |
| $T_7$ | 10.072 | 35.347 | 2e-09 |
| $R_4$ | -4.002 | 5.430 | 0.020 |
| $R_5$ | -4.029 | 15.494 | 8e-05 |
| $R_6$ | -8.047 | 10.067 | 0.002 |

The model performance was evaluated using leave-one-out cross-validation (LOOCV) and leave-one-year-out (also called leave-three-out) cross-validation (LOYOCV). For LOOCV, the observation in one area in a given year was removed before refitting the model. The best-fitting model successfully predicted the major outbreaks by year and by area using LOOCV (Fig. 5a-c, supplementary Table S2 and supplementary dataset S1 online). 42 out of a total of 51 (82.4%) total observations by year and by area (annual incidence rates in 17 years and 3 areas) were within the 95% confidence interval of the annual incidence predicted by our model. Note that the upper and the lower bounds of the confidence intervals were rounded to the nearest integer when assessing the prediction accuracy of observations in Table S2. 5 out of 6 observations of area-specific outbreaks in 2002 and 2018 were able to be predicted. Although the 2018 outbreak in NTS was not predicted, the model still identified an outbreak (defined as the annual number of dengue cases *>* 2) in that area. When LOYOCV was used, the model predicted outbreaks (defined as the annual number of dengue cases *>* 2) in 5 years (i.e., the years 2002, 2003, 2007, 2015 and 2018), which include the two major outbreak years (2002 and 2018) and a year with sporadic cases (2015) (Fig. 5d).

**Figure 5.**
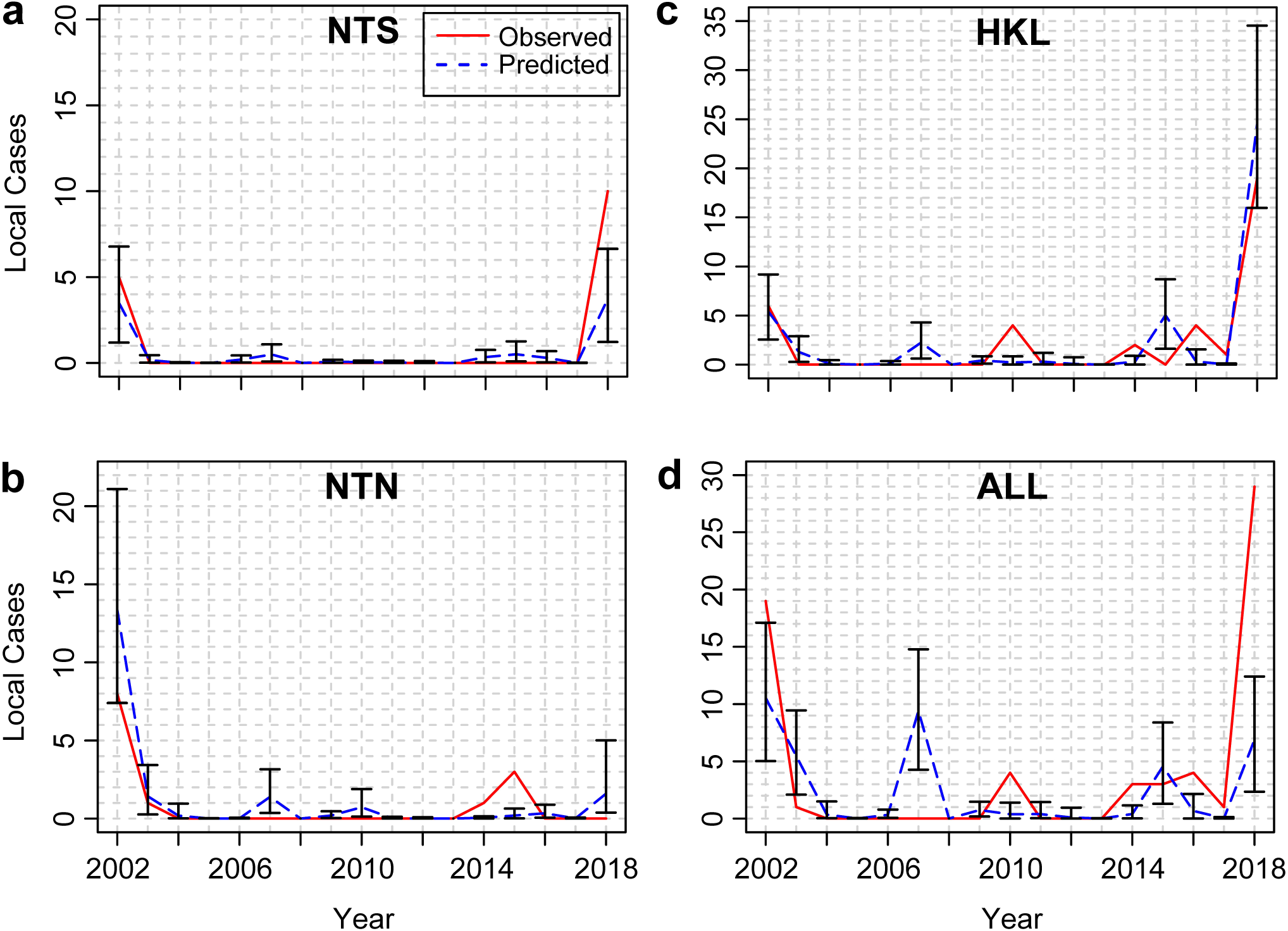
Comparison between observed and predicted number of annual dengue cases. (a-c) Observed and predicted number of annual dengue cases in each pre-defined area (NTS, NTN, and HKL) with 95% confidence intervals using leave-one-out cross-validation. The predicted values represent the mean of the Poisson distribution of the annual dengue cases in each area, estimated using a generalized linear mixed model with a restricted maximum likelihood method. (d) Observed and predicted number of annual dengue cases in the whole Hong Kong area (ALL) with 95% confidence intervals using leave-one-year-out cross-validation.

The best-fitting model produced an *MSE*_*tr*_ of 0.592, an *MSE*_*va*_ of 3.538 and an *MSE*_*ratio*_ (*MSE*_*va*_*/MSE*_*tr*_) of 5.976 using LOOCV (Table 3). Both the *MSE*_*va*_ and the *MSE*_*ratio*_ of the best-fitting model were the lowest compared with the alternative (Model: *T*_*mean*_+*R*_*max*_), fixed effects (same predictors as the best-fitting model but without random effects), and full models, indicating the best prediction performance among all the tested models. The alternative model using *T*_*mean*_ and *R*_*max*_ produced an *MSE*_*tr*_ of 0.672 and an *MSE*_*va*_ of 5.143, which confirmed that the chosen best model fitted the data better than the alternative. The fixed effects model performed similarly to but slightly worse than the best-fitting model. The full model, including all predictors, had the lowest *MSE*_*tr*_ value but the highest *MSE*_*va*_ and the highest *MSE*_*ratio*_, indicating an overfitting phenomenon. The best model also performed better than the other models when the Normalised Mean Squared Error (NMSE) was used. When LOYOCV was used, the *MSE*_*ratio*_ (12.629) of the best-fitting model was slightly higher than that obtained from LOOCV results. Although the fixed effects model can perform similarly or slightly better according to the *MSE*_*va*_ (or *NMSE*_*va*_), the best-fitting model performed better than the other models according to the *MSE*_*ratio*_ (or *NMSE*_*ratio*_) (Supplementary Table S3). These results demonstrate that while working with a small number of dengue observations in Hong Kong, a mixed effects model with AICc-selected variables can reduce model overfitting.

**Table 3.** The mean squared errors (MSE) and normalised mean squared errors (NMSE) of each model tested using leave-one-out cross-validation. The MSE and NMSE for the training and the validation sets, and the ratios of the MSE and NMSE of the validation set to that of the training set, respectively, are listed. The best-fitting model (Model: *T*_*mean*_ + *R*_*tot*_) was compared with the alternative model (Model: *T*_*mean*_ + *R*_*max*_), the fixed effects model (same predictors as the best-fitting model but without random effects) and the full model (mixed effects with all predictors).

|  | MSE |  |  |  | NMSE |  |  |
| --- | --- | --- | --- | --- | --- | --- | --- |
|  | *Training* | *Validation* | *Ratio* |  | *Training* | *Validation* | *Ratio* |
| Best-fitting | 0.592 | 3.538 | 5.976 |  | 0.032 | 0.186 | 5.813 |
| Alternative | 0.672 | 5.143 | 7.653 |  | 0.036 | 0.271 | 7.528 |
| Fixed effects | 0.551 | 3.543 | 6.430 |  | 0.029 | 0.186 | 6.414 |
| Full | 0.408 | 189.393 | 464.199 |  | 0.022 | 9.968 | 453.091 |

### Effects of Climate Variables

The predictive model showed that all monthly mean temperature predictors (*T*_3_, *T*_4_, *T*_5_, and *T*_7_) were positively correlated with the number of annual cases, while monthly total rainfall predictors (*R*_4_, *R*_5_, and *R*_6_) were all negatively correlated. These behaviors were also found in the stepAICc results of the other 3 models with different climate parameter sets (Supplementary Table S1), confirming the positive correlations of the temperature and the negative correlations of the early rainfall predictors with the number of annual cases. Among all the temperature predictors, *T*_7_ (the mean temperature in July) was the most significant predictor (*p <* 10^−8^) and its regression coefficient had the highest magnitude, indicating the strongest effect. The coefficients of the 3 rainfall predictors (from April to June) were of similar magnitude, indicating a much longer delayed effect of rainfall compared to temperature. In the NTS and HKL areas, relatively high values for *T*_3_, *T*_5_, and *T*_7_ and relatively low values for *R*_4_ and *R*_5_ were found in 2018, which may explain the the outbreak in 2018 in those two areas.

We further examined the marginal effects of the 4 predictors (*T*_7_, *R*_4_, *R*_5_, and *R*_6_) with the greatest magnitude of regression coefficients on the relative risk *RR* (Supplementary Fig. S1). *RR* was defined as the number of annual cases divided by the average number of annual cases. Marginal effects of the collected temperature (without normalization) 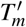 and rainfall 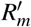 (without normalization) were also obtained after refitting the same model (Supplementary Fig. S2). If the rainfall is lower than 200mm (per month) between May and June, or lower than 100mm in April, a higher relative risk of dengue incidence is observed.

## Discussion

This study is the first to explore the association between seasonal climate variables and dengue incidence while taking into account the spatial heterogeneity of the dengue cases in Hong Kong. After dividing Hong Kong into 3 areas, a Poisson mixed effects model was fitted using the monthly temperature and rainfall data from the early months of the year to forecast annual dengue incidence in the defined areas. Given that most of the dengue infections occurred in August and September, the model successfully predicts annual dengue incidence using seasonal weather data from the preceding months. Instead of using a distributed lag model, our approach allows us to capture the impact of the weather in each month on the annual number of dengue cases, which is more suitable for regions affected by seasonal epidemics^8^.

Because most of the outbreaks occurred in August and September, the period of the delay of climate effects can be estimated. Our results showed that the temperature is positively associated with the relative risk within time lags of at least 1 or 2 months (from *T*_7_ to the start of the outbreak), which is consistent with previous studies^17, 30–34^. These lagged effects could be caused by the incubation periods of dengue viruses and the mosquito’s reproductive life cycle. The delayed effect of rainfall from between 3 and 5 months (from *R*_4_, *R*_5_ and *R*_6_ to the start of the outbreak) in our study is also consistent with recent findings that rainfall can be negatively associated with the risk of dengue outbreaks with a lead time up to 5 months^17^.

The three defined areas NTS, NTN, and HKL in our study were selected to reflect the spatial heterogeneity of the dengue cases in Hong Kong. This spatial heterogeneity may be caused by area-specific factors other than climate variables; for instance, urbanisation, transportation, population density, etc., could all contribute to the expansion of dengue. For the purpose of our study, as we mainly focus on the impact of climate, those area-specific factors were considered as random effects in our model. Introducing random effects into the model can lead to higher accuracy of parameter estimation especially when the dataset contains non-independent observational units that are hierarchical in nature^35^.

The significance of understanding the climate-dengue association lies in three aspects. First, being a highly populated region with more than 50 million people travelling to the city annually^36^, infectious diseases can spread rapidly not only within Hong Kong but also to other countries due to its high level of contact mixing within- and between-countries^37–39^. Therefore, whether an even larger dengue outbreak will occur in Hong Kong in the future has become a primary global health concern.

Second, we demonstrated that in addition to global warming, the variation in rainfall intensity associated with the East Asian monsoon may play a significant role in the recent dengue expansions in some East Asian subtropical countries/cities. An increasing risk of seasonal dengue epidemics has been observed in both southern Taiwan and Hong Kong in the recent years, and similar to our findings, a negative association between early rainfall and dengue infections has previously been identified in southern Taiwan^8^. As Hong Kong and Taiwan are both located in the East Asian subtropics, they are both affected by the East Asian rainy season occurring annually between February and June^40^. Our results suggest that the variability during or even before the East Asian monsoon (pre-summer) rainy season may pose a high risk for dengue expansion in the region and complex non-linear effects should be studied further. As for Guangzhou, we noticed that before the 2014 dengue outbreak period (the largest one in Guangzhou), heavy rainfall mixed with drought (or low initial water level) were observed in that particular year^41, 42^.

Third, our findings on rainfall effects can be used to inform mosquito control strategies in Hong Kong and possibly in other nearby countries similarly affected by the East Asian monsoon. The authorities of Hong Kong often enhance mosquito control measures while increasing public awareness especially during periods of heavy rainfall^43, 44^. Our results suggest that in this particular region, the relative risk of incidence is higher when the intensity of rainfall before summer is lower. This implies that stepping up mosquito control measures to reduce mosquito activities is the most critical when the early rainfall amount is lower than the expected level.

In conclusion, we have developed a generalized linear mixed model that is able to successfully forecast annual dengue incidence across Hong Kong by taking into account both the climate data and the unobserved area-specific factors. Our findings may provide a valuable assessment on how to plan dengue prevention and control measures more effectively in Hong Kong.

## Methods

### Study Area

Hong Kong is officially divided into 18 districts, which are grouped into three main areas: Hong Kong Island, Kowloon, and the New Territories (Supplementary Fig. S3). Based on the geography, population density, and the usage of the Mass Transit Railways (MTR), which is the most heavily used public transportation in Hong Kong, we redivided the city into three main areas. We first divided the New Territories into two regions: New Territories-South (NTS) and New Territories-North (NTN) such that NTS consists of the smaller and outlying islands while NTN consists of the region connected to Mainland China. We then merged Hong Kong Island and Kowloon into Hong Kong Island & Kowloon (HKL) because of their high similarities in terms of population density (Supplementary Fig. S4) and public transportation network. Kwai Tsing District was reassigned from the New Territories into this area as it also has a high population density, is located close to Kowloon, and is linked to the area by the busiest MTR line^45^.

### Dengue Data

The confirmed numbers of local dengue fever cases from 2002 to 2018 were collected through press releases issued by the Centre for Health Protection (CHP) of the Department of Health of the Hong Kong Government^46^. The official statements indicate the location and the date of every local case confirmed through laboratory tests such as NS1 antigen and IgM serological assays, where the possible infection location is determined by either the residence or the travel history of the patient. The annual number of dengue cases was defined as the total number of confirmed local (indigenous) cases in each year (Fig. 1 inset); the onset of dengue fever symptoms in all years between 2002 and 2018 was observed between June and December (Fig. 3). Imported cases, confirmed by CHP, were excluded in order to rule out travel-related factors influencing infection rates.

### Meteorological Data

Both the daily mean temperature and the daily total rainfall from 2002 to 2018 inclusive were retrieved from the Hong Kong Observatory^47^. A total of 11 out of the available 49 automatic weather stations were selected to represent all districts in Hong Kong (Fig. 1). The stations were chosen as they provided the most complete temperature and rainfall data in those years; any weather stations with more than 30% missing data were excluded. The selected station with the highest number of missing data is in Tuen Mun District since it only started recording from 2007 onwards. Two additional automatic rainfall stations on the island were selected due to the incomplete rainfall data from the weather stations in Hong Kong Island. Daily weather data were then averaged across different stations in each defined area to obtain average area-specific weather data. Monthly mean (minimum) temperature is defined as the average (minimum) of the daily mean temperature while the monthly total (maximum) rainfall is defined as the total (maximum) of the daily total rainfall. In general, a strong seasonal pattern was observed among the climate parameters from 2002 to 2018 (Fig. 2). An increasing trend in monthly temperature and rainfall was observed from January until July and June, respectively.

### Model Framework

Given the similar seasonality of dengue epidemics in Southern Taiwan and Hong Kong where infections mostly appeared in the latter half of the year starting in summer, we extended a previous study of forecasting dengue annual cases in Southern Taiwan by fitting a model using the temperature and rainfall data from the earlier part of the year to predict the subsequent area-specific outbreaks^8^. In order to account for any unobserved factors relating to the studied areas, we developed a Poisson generalized linear mixed model (GLMM) to fit the confirmed cases in the different defined areas. Let 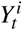 denote the annual number of dengue cases and 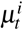 denote the expected number of dengue cases, which is the corresponding distribution mean in area *i* and year *t*, then:

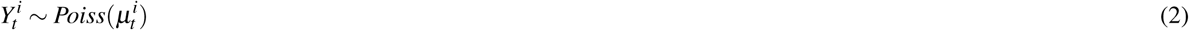

and the model is specified by:

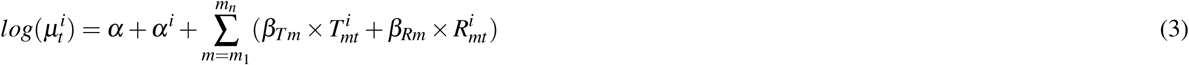

where 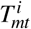 and 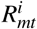 denote the *m*^*th*^ month’s (ranges from *m*_1_ to *m*_*n*_) normalised temperature and normalised rainfall predictors in the year *t* and area *i, β*_*Tm*_ and *β*_*Rm*_ denote the regression coefficient of 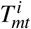 and 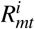 respectively, *α* represents the intercept, and *α*^*i*^ represents the area-specific random intercept.

The monthly predictors were obtained using monthly climate data that were normalised into the range [0,1] from the collected temperature 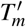 and rainfall 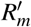 data, so that the effects of the predictors on the responses are easier to compare when fitting the model^33, 48^:

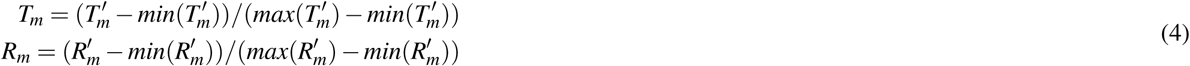

Despite a number of zero counts in the dengue case data for certain years, we did not use zero-inflated models as they assume that excess zeros are generated by a separate process from the count values (e.g. underreporting of infections). We assumed that the zero values represent the true case numbers as CHP conducts strict dengue prevention and control measures in Hong Kong.

### Model Selection

A two-step procedure was used for the model selection. First, a candidate model was chosen to represent each of the four climate parameter sets (*T*_*mean*_ or *T*_*min*_ with *R*_*tot*_ or *R*_*max*_). The stepwise algorithm based on the Akaike Information Criterion with correction (AICc), also known as the stepAICc, was used to select the fixed effect variables (monthly climate predictors) for each of the candidate models in a forward selection approach (Supplementary Fig. S5). The stepwise algorithm has been commonly used in selecting relevant environmental or climatic determinants to predict disease occurrences^8, 30^. This approach iteratively added a new variable to the model, starting with zero variables, to fulfil the selection criteria (such as smaller AIC or AICc) until the most optimal set of variables was reached. For our case, the AICc was used instead of AIC due to the small number of observations. For the purpose of model selection, all the models were estimated using Maximum Likelihood by Laplace approximation provided by the R package, glmmTMB^49^, in order to compare models with different fixed effects. Laplace approximation is a suggested method for fitting Poisson mixed effects models^50^. Secondly, the candidate models with the most optimal set of variables were then compared with each other based on their AICc and Bayesian Information Criterion (BIC). The model with the lowest AICc and BIC (or only AICc) was chosen as the best-fitting model.

### Model Evaluation

Leave-one-out cross-validation (LOOCV) was used in order to evaluate the best-fitting model’s overall performance and to ensure that the model does not overfit. During each iteration, the observation of the annual number of dengue cases in a particular year and area was excluded while fitting the model using the restricted maximum likelihood method. The number of annual dengue cases of this particular observation was then estimated (predicted) given the coefficients produced by the fitted model in that iteration. In order to measure the performance of the model against the observation data, the Mean Squared Error (MSE) was calculated for both training and validation sets using the following general formulae:

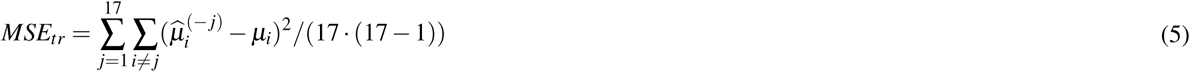

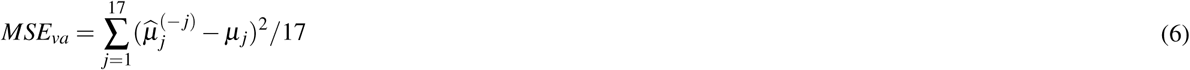

where *j* indicates the index of a year to be tested, 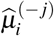 indicates the estimated number of cases in year *i* when the number in year *j* was removed, and *µ*^*i*^ is the observed number of cases in year *i*. When pre-defined areas were considered, the highest value of index *j* became 51 to represent the total number of observations of annual dengue incidence by year and by area. In addition, leave-one-year-out (leave-three-out) cross-validation (LOYOCV) was also used to evaluate the best-fitting model when the observations of the annual number of dengue cases in all three areas in a particular year were removed and used as the validation set while fitting the model.

Bootstrapped confidence intervals were computed. For each predicted 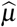 of a model, we generated 1000 samples of outcomes from a Poisson distribution with the rate parameter equal to the estimated parameter 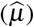 of that fitted model. Based on these 1000 simulated outcomes, we refitted the model 1000 times to obtain the distribution of the parameter’s values. The 95% confidence interval defines a range of values between the 97.5% and 2.5% quantiles. We considered the model to be satisfactory if it managed to successfully predict the two major outbreaks (within confidence intervals) in 2002 and 2018 in the LOOCV process and if it produced relatively low values for *MSE*_*va*_ or *MSE*_*ratio*_ (defined as *MSE*_*va*_*/MSE*_*tr*_). The Normalised Mean Squared Error (NMSE) was also calculated by dividing the MSE of the validation or training set by the range of the observed variable.

## Supporting information

Suplplementary Data Set

## Acknowledgements

We would like to thank Drs. Wen Zhou (City University of Hong Kong), Xiaolin Zhu (Hong Kong Polytechnic University) and James Hay (Imperial College London) for their valuable comments on model development, weather patterns and geographical information. We also thank the financial supports from startup grant and the Applied Strategic Development Centres at City University of Hong Kong.

## Author contributions statement

K.S. and M.M.T. conducted data collection. K.S., J.L., M.M.T., P.L. and H.Y. conducted the statistical model development and wrote the main manuscript text, K.S. and H.Y. analysed the results. All authors reviewed the manuscript.

## Additional information

## Data Availability

The climate data are available on the Hong Kong Observatory website while the dengue cases data are available on the online Press Release published by the Centre for Health Protection, Hong Kong. The overall dataset including reformatted climate data and the R code files produced and applied in this study are available from the corresponding author on request.

