## Supplementary Figures and Tables for "The effects of seasonal climate variability on dengue annual incidence in Hong Kong: A modelling study"

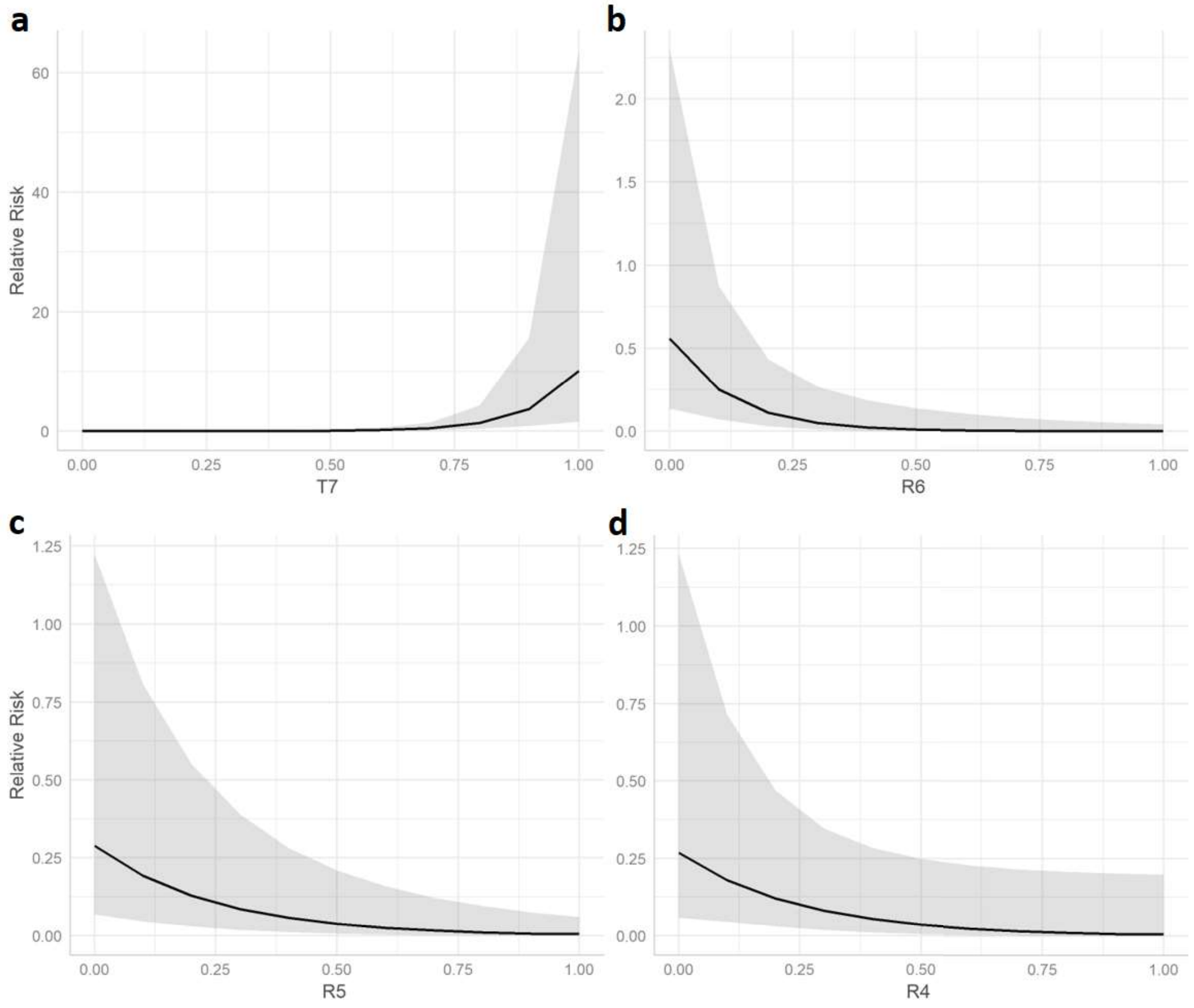


**Figure S1.** The marginal effects of the 4 predictors with the greatest magnitude of correlation coefficients on the relative risk (RR). The RR is defined as the predicted number of annual cases divided by the average number of annual cases over all years from 2002 to 2018. The marginal effects of (a) $T_7$, (b) $R_6$, (c) $R_5$ and (d) $R_4$ are included.

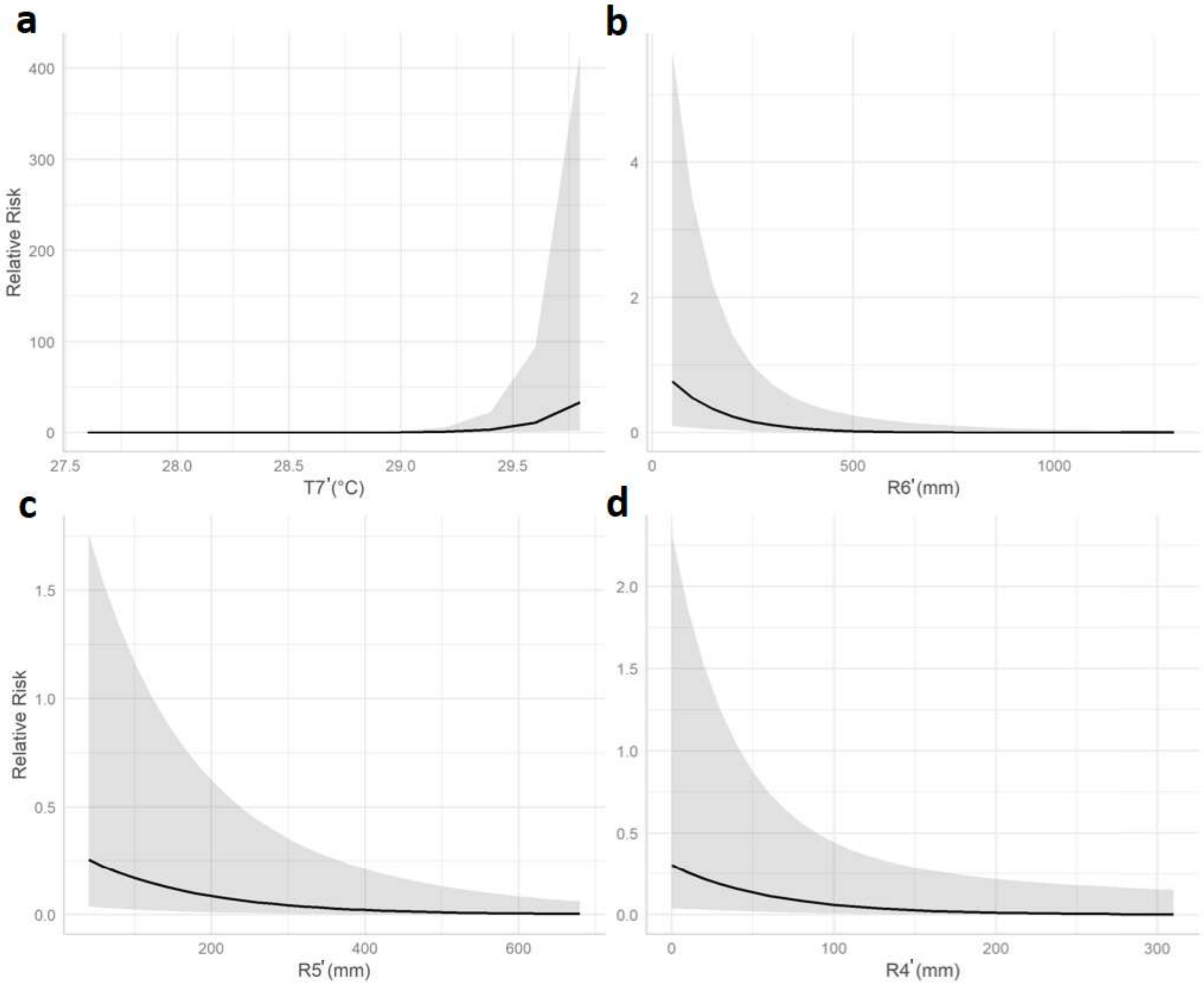


**Figure S2.** The marginal effects of the 4 predictors, using the original weather data, with the greatest magnitude of correlation coefficients on the relative risk (RR). The RR is defined as the number of annual cases divided by the average number of annual cases over all years from 2002 to 2018. The marginal effects of (a) $T_7'$, (b) $R_6'$, (c) $R_5'$ and (d) $R_4'$ are included. Note that the range of each predictor is in the original units but not the standardised ratio for easier comparison with the actual monthly climate data.

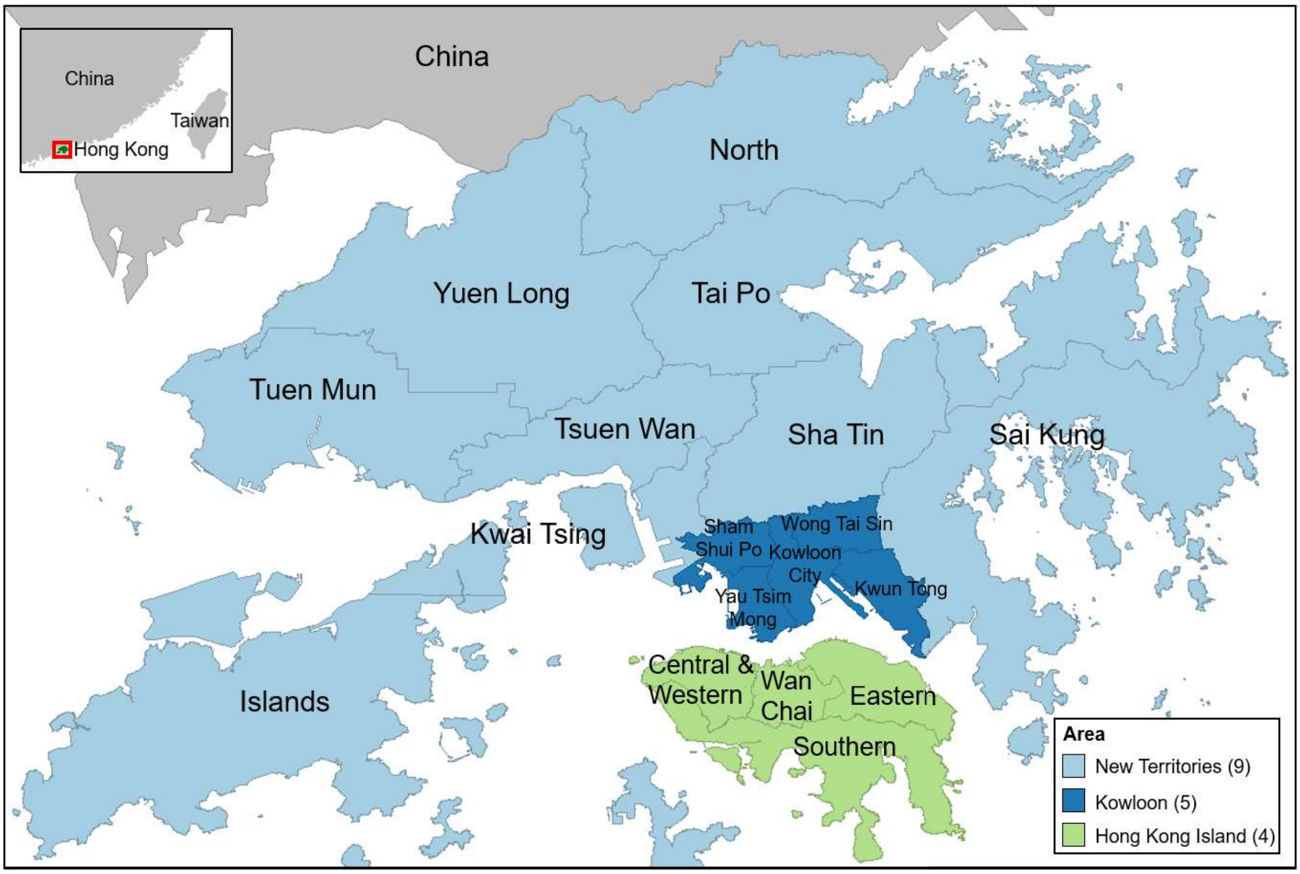


**Figure S3.** The official districts and regions of Hong Kong. Numbers in the legends indicate the number of districts belonging to each region.

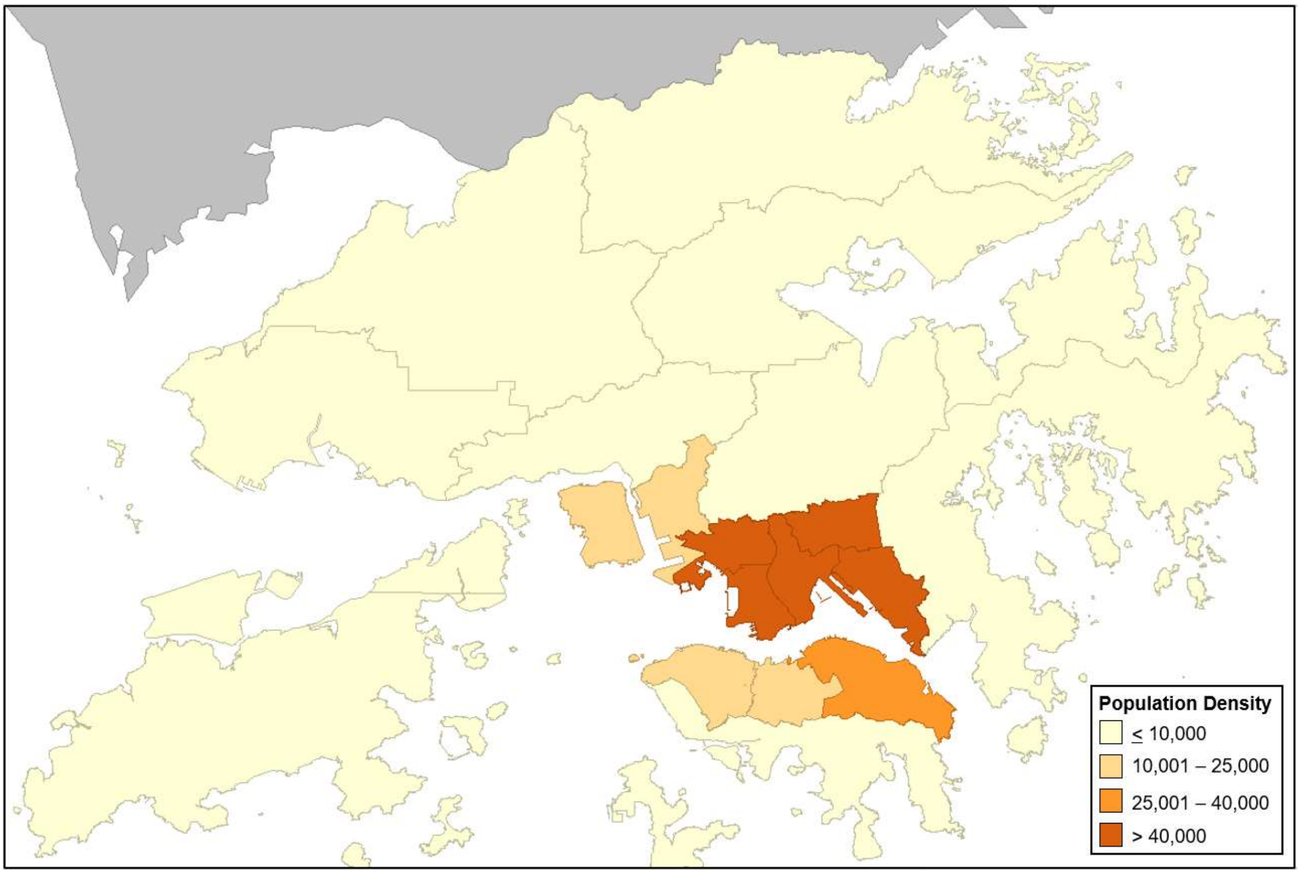


**Figure S4.** Hong Kong population density map. Density is defined as the total number of population divided by area in $km^2$.

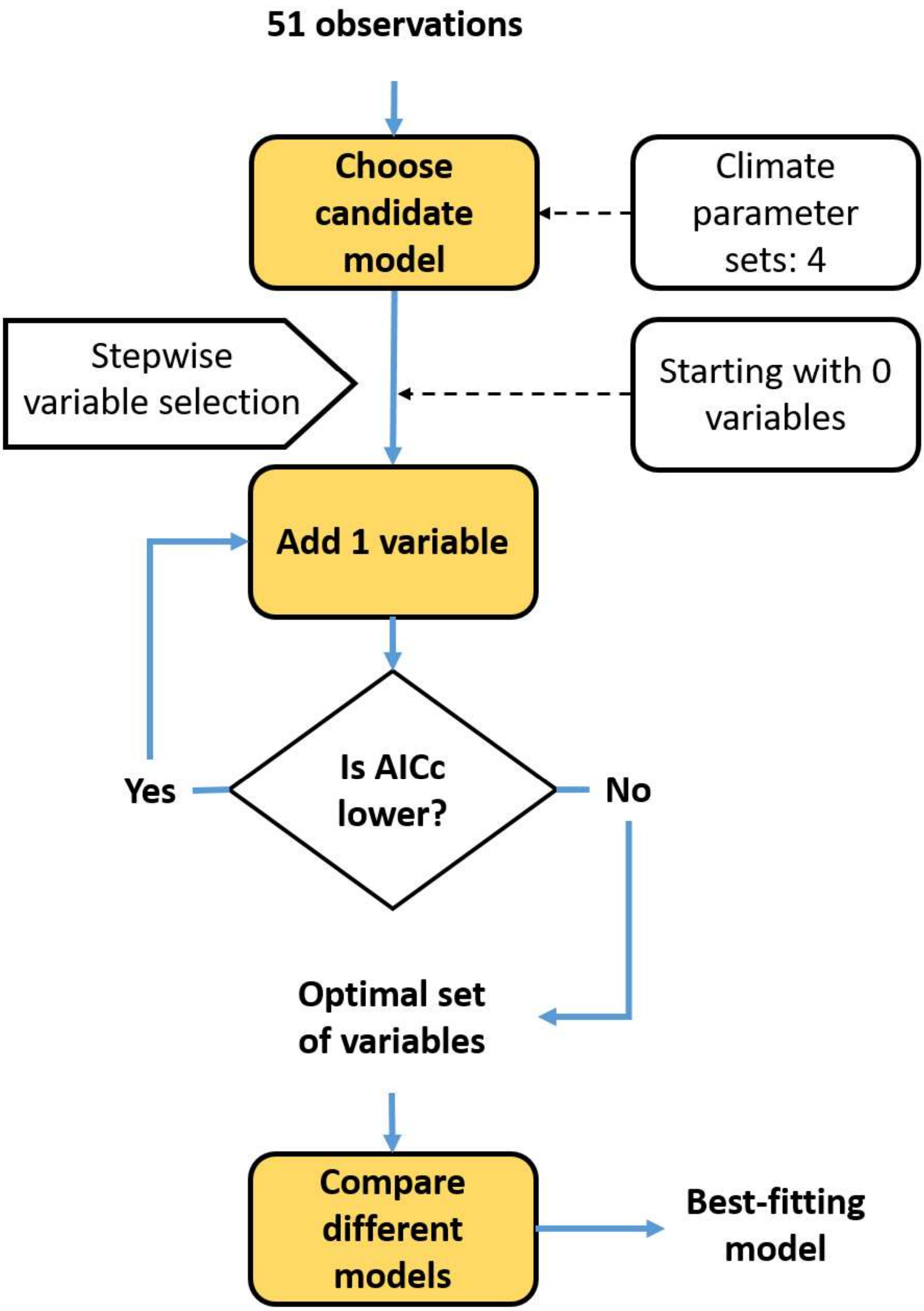


**Figure S5.** Flowchart for selecting the best-fitting model using a two-step procedure. The procedure involves variable selection for a candidate model with a climate parameter set as the first step and model comparison between four different climate parameter sets as the second step. The initial data includes a total of 51 observations in the pre-defined areas. The stepwise algorithm based on the Akaike Information Criterion with a correction (AICc) is used to produce the optimal set of variables.

**Table S1.** The variable selection stages of the stepwise forward selection algorithm with AICc for each set of monthly climate predictors. $T_m$ and $R_m$ denote the temperature and rainfall predictors in the $m^{th}$ month of the year. The order of the rows indicates the AICc results at each selection step. The order of the columns (from the left to right) represents the sequence of the variables that have been selected. A newly selected variable is listed to the right of the last variable. The optimal set of variables is listed for each climate parameter combination set (as in subtable i to iv).

(i) Monthly mean temperature ($T_{mean}$) and monthly total rainfall ($R_{tot}$)

| Variables | | | | | | | | *AICc* |
|---|---|---|---|---|---|---|---|---|
| 0 | $T_5$ | $T_3$ | $T_7$ | $T_4$ | $R_5$ | $R_6$ | $R_4$ | |
| 0.138 | | | | | | | | 266.79 |
| -2.531 | 4.858 | | | | | | | 169.03 |
| -4.732 | 3.454 | 4.372 | | | | | | 139.49 |
| -7.019 | 2.982 | 5.827 | 2.807 | | | | | 130.95 |
| -10.903 | 2.475 | 6.070 | 6.043 | 3.323 | | | | 116.78 |
| -10.626 | 0.670 | 6.284 | 7.366 | 4.325 | -2.311 | | | 113.31 |
| -9.172 | 1.543 | 5.852 | 8.273 | 3.222 | -3.292 | -5.158 | | 110.30 |
| -6.200 | 1.210 | 3.594 | 10.072 | 2.542 | -4.029 | -8.047 | -4.002 | 107.84 |

(ii) Monthly mean temperature ($T_{mean}$) and monthly maximum rainfall ($R_{max}$)

| Variables | | | | | | | | *AICc* |
|---|---|---|---|---|---|---|---|---|
| 0 | $T_5$ | $T_3$ | $T_7$ | $T_4$ | $R_5$ | $R_4$ | $R_8$ | |
| 0.138 | | | | | | | | 266.79 |
| -2.531 | 4.858 | | | | | | | 169.03 |
| -4.732 | 3.454 | 4.372 | | | | | | 139.49 |
| -7.019 | 2.982 | 5.827 | 2.807 | | | | | 130.95 |
| -10.903 | 2.475 | 6.070 | 6.043 | 3.323 | | | | 116.78 |
| -9.751 | 1.644 | 5.315 | 6.438 | 3.578 | -2.051 | | | 114.70 |
| -6.544 | 0.873 | 3.406 | 6.860 | 2.720 | -2.719 | -5.613 | | 110.93 |
| -5.589 | 0.302 | 2.943 | 7.608 | 3.551 | -3.787 | -6.803 | -2.789 | 108.84 |

(iii) Monthly minimum temperature ($T_{min}$) and monthly total rainfall ($R_{tot}$)

| Variables | | | | | | | | *AICc* |
|---|---|---|---|---|---|---|---|---|
| 0 | $R_4$ | $R_5$ | $T_5$ | $T_6$ | $T_7$ | $R_6$ | $R_8$ | |
| 0.138 | | | | | | | | 266.79 |
| 1.549 | -6.125 | | | | | | | 203.50 |
| 2.334 | -5.657 | -2.827 | | | | | | 184.77 |
| -0.435 | -4.271 | -3.509 | 3.927 | | | | | 175.82 |
| -2.720 | -3.467 | -3.245 | 3.963 | 2.843 | | | | 167.39 |
| -4.726 | -3.370 | -3.234 | 4.748 | 2.998 | 2.393 | | | 160.83 |
| -2.646 | -3.831 | -6.277 | 4.169 | 2.068 | 5.501 | -7.796 | | 145.04 |
| -1.368 | -5.076 | -3.238 | 1.130 | 0.540 | 5.077 | -9.231 | 4.109 | 134.34 |

(iv) Monthly minimum temperature ($T_{min}$) and monthly maximum rainfall ($R_{max}$)

| Variables | | | | | | | | | | *AICc* |
|---|---|---|---|---|---|---|---|---|---|---|
| 0 | $R_4$ | $R_5$ | $T_7$ | $R_6$ | $R_8$ | $T_5$ | $T_8$ | $R_3$ | $T_4$ | |
| 0.138 | | | | | | | | | | 266.79 |
| 1.640 | -9.323 | | | | | | | | | 205.61 |
| 2.378 | -6.652 | -4.518 | | | | | | | | 169.47 |
| 0.159 | -10.200 | -4.894 | 4.993 | | | | | | | 143.309 |
| 0.779 | -10.306 | -6.008 | 6.708 | -5.135 | | | | | | 132.57 |
| 1.501 | -10.529 | -6.244 | 6.450 | -4.775 | -1.861 | | | | | 130.05 |
| -0.070 | -9.798 | -5.949 | 6.592 | -4.332 | -2.235 | 1.928 | | | | 129.90 |
| 0.695 | -10.564 | -6.385 | 7.791 | -5.130 | -3.469 | 2.774 | -2.242 | | | 127.43 |
| -2.322 | -10.738 | -6.678 | 9.695 | -1.348 | -6.416 | 6.404 | -4.650 | 3.277 | | 118.64 |
| -3.322 | -10.563 | -6.598 | 9.023 | -0.552 | -6.632 | 7.145 | -5.510 | 2.882 | 2.275 | 117.91 |

**Table S2.** The observed and predicted results with 95% confidence intervals in the defined areas using leave-one-out cross-validation. Note that the predicted numbers and upper and lower bounds of confidence intervals were rounded to the nearest integer.

| **NTS** | 2002 | 2003 | 2004 | 2005 | 2006 | 2007 | 2008 | 2009 | 2010 | 2011 | 2012 | 2013 | 2014 | 2015 | 2016 | 2017 | 2018 |
|---|---|---|---|---|---|---|---|---|---|---|---|---|---|---|---|---|---|
| Observed | 5 | 0 | 0 | 0 | 0 | 0 | 0 | 0 | 0 | 0 | 0 | 0 | 0 | 0 | 0 | 0 | 10 |
| Predicted | 3 | 0 | 0 | 0 | 0 | 0 | 0 | 0 | 0 | 0 | 0 | 0 | 0 | 1 | 0 | 0 | 4 |
| CI (lower) | 1 | 0 | 0 | 0 | 0 | 0 | 0 | 0 | 0 | 0 | 0 | 0 | 0 | 0 | 0 | 0 | 1 |
| CI (upper) | 7 | 0 | 0 | 0 | 0 | 1 | 0 | 0 | 0 | 0 | 0 | 0 | 1 | 1 | 1 | 0 | 7 |

| **NTN** | 2002 | 2003 | 2004 | 2005 | 2006 | 2007 | 2008 | 2009 | 2010 | 2011 | 2012 | 2013 | 2014 | 2015 | 2016 | 2017 | 2018 |
|---|---|---|---|---|---|---|---|---|---|---|---|---|---|---|---|---|---|
| Observed | 8 | 1 | 0 | 0 | 0 | 0 | 0 | 0 | 0 | 0 | 0 | 0 | 1 | 3 | 0 | 0 | 0 |
| Predicted | 13 | 1 | 0 | 0 | 0 | 1 | 0 | 0 | 1 | 0 | 0 | 0 | 0 | 0 | 0 | 0 | 2 |
| CI (lower) | 7 | 0 | 0 | 0 | 0 | 0 | 0 | 0 | 0 | 0 | 0 | 0 | 0 | 0 | 0 | 0 | 0 |
| CI (upper) | 21 | 3 | 1 | 0 | 0 | 3 | 0 | 0 | 2 | 0 | 0 | 0 | 0 | 1 | 1 | 0 | 5 |

| **HKL** | 2002 | 2003 | 2004 | 2005 | 2006 | 2007 | 2008 | 2009 | 2010 | 2011 | 2012 | 2013 | 2014 | 2015 | 2016 | 2017 | 2018 |
|---|---|---|---|---|---|---|---|---|---|---|---|---|---|---|---|---|---|
| Observed | 6 | 0 | 0 | 0 | 0 | 0 | 0 | 0 | 4 | 0 | 0 | 0 | 2 | 0 | 4 | 1 | 19 |
| Predicted | 5 | 1 | 0 | 0 | 0 | 2 | 0 | 0 | 0 | 0 | 0 | 0 | 0 | 5 | 0 | 0 | 25 |
| CI (lower) | 3 | 0 | 0 | 0 | 0 | 1 | 0 | 0 | 0 | 0 | 0 | 0 | 0 | 2 | 0 | 0 | 16 |
| CI (upper) | 9 | 3 | 0 | 0 | 0 | 4 | 0 | 1 | 1 | 1 | 1 | 0 | 1 | 9 | 2 | 0 | 35 |

**Table S3.** The mean squared errors (MSE) and normalised mean squared errors (NMSE) of each model tested using leave-one-year-out (leave-three-out) cross-validation. The MSE and NMSE for the training and the validation sets, and the ratios of the MSE and NMSE of the validation set to that of the training set, respectively, are listed. The best-fitting model was compared with the alternative model (Model: $T_{mean} + R_{tot}$), the fixed effects model (same predictors as the best-fitting model but without random effects) and the full model (all predictors).

| | MSE | | | NMSE | | |
|---|---|---|---|---|---|---|
| | *Training* | *Validation* | *Ratio* | *Training* | *Validation* | *Ratio* |
| Best-fitting | 0.591 | 7.464 | 12.629 | 0.033 | 0.393 | 11.559 |
| Alternative | 0.650 | 11.603 | 17.851 | 0.037 | 0.611 | 16.514 |
| Fixed effects | 0.547 | 7.291 | 13.329 | 0.031 | 0.384 | 12.387 |
| Full | 0.366 | >1e04 | >1e04 | 0.021 | >1e04 | >1e04 |
